# Lineage-specific evolution of regulatory landscapes in a polyploid plant and its diploid progenitors

**DOI:** 10.64898/2025.12.04.692133

**Authors:** Xiang Li, Xuan Zhang, Ziliang Luo, Hao Zhang, John Pablo Mendieta, Robert J. Schmitz

## Abstract

*Cis*-regulatory elements are specific DNA sequences that control gene expression in a spatiotemporal manner, and variation within these elements represents a major source of phenotypic diversity and evolutionary innovation. Nevertheless, how regulatory elements evolve and shape gene expression remains poorly understood, particularly in plants. The well-resolved phylogeny of allopolyploid peanut (*Arachis hypogaea*) and its diploid progenitors, *A. duranensis* and *A. ipaensis*, provides an ideal system to investigate the regulatory evolution at a lineage-specific level. By integrating comparative analyses of sequence similarity, chromatin accessibility, histone modifications, conserved noncoding sequences (CNSs), and gene expression, we reconstructed the evolutionary trajectories of Accessible Chromatin Regions (ACRs), where regulatory elements typically reside, and revealed their distinct contributions to homeolog expression bias, unequal expressions between homeologs. Most ACRs exhibited high sequence similarity, comparable chromatin accessibility, and conserved states for H3K4me3, H3K56ac, and H3K36me3, indicating regulatory stability after hybridization and polyploidization. However, a subset of novel ACRs emerged *de novo* from previously non-regulatory regions or through sequence mutations in preexisting ACRs, arising at different rates and evolutionary stages. Notably, even highly sequence-conserved ACRs exhibited substantial variation in chromatin accessibility, consistent with CNS composition differences and minor sequence variation, although causal relationships remain to be demonstrated. Our analyses further revealed a complex spectrum of CNS dynamics across the diploid-polyploid framework. Overall, our study provides empirical insights into the fine-scale evolution of plant regulatory landscapes and complements previous large-scale comparisons across distant lineages.

## Introduction

*Cis*-regulatory elements, specific DNA sequences bound by transcription factors, regulate gene expression in a spatiotemporal manner (1–4). Genetic variation within these elements represents an important source of phenotypic diversity and evolutionary innovation (5, 6). Nevertheless, the origin and the extent of sequence divergence within these elements and how such divergence shapes regulatory activity and gene expression remain poorly understood, especially in plants.

Unlike the relatively conserved and stable mammalian genomes, plant genomes are highly dynamic, exhibiting extensive variation in size, structure, and compaction (7, 8). Frequent macro- and micro-genomic rearrangements, whole genome duplication, gene duplication and unequal gene retention have all shaped plant genome architecture. These processes likely exert a profound impact on the emergence and diversification of regulatory sequences (9–11). Comparative analyses across distantly related plants have revealed a high rate of turnover in regulatory regions, underscoring the fast-evolving nature of plant *cis*-regulatory landscapes and the difficulty of tracing their evolutionary trajectories (12–14). Yet, deeply conserved non-coding regions (CNSs) spanning more than 280 plant species and 300 million years of diversification have also been identified (10). To reconcile this seemingly contradictory observation between, rapid turnover versus deep conservation, and to complement broad-scale comparative studies, analyses among closely related species provides a critical and trackable framework to uncover how *cis*-regulatory sequences diversify, acquire new functions, and drive regulatory innovation over short evolutionary timescales.

Polyploid plants that have undergone whole-genome duplication provide a natural context to contrast the evolutionary fates of duplicated regulatory elements. During polyploidization, regulatory elements are copied alongside genes and may experience relaxed selective constraints like those acting on duplicated coding sequences (15), thereby enabling divergence in both sequence and function (5). Recent studies in soybean, cotton, and strawberry have illustrated that variation in *cis*-regulatory regions contributes to differential expression among duplicated genes (16–22). However, most insights come from species with ancient duplication events or from analyses lacking direct comparisons with their progenitors. Incorporating progenitor species with polyploid species provides a unique opportunity to reconstruct how regulatory sequences originate and diversify.

This framework is particularly well suited for recently formed allopolyploids such as cultivated peanut (*Arachis hypogaea*), an allotetraploid crop (AABB, 2n = 4x =40) derived from hybridization between two diploid progenitors, *A. duranensis* (progenitor A) and *A. ipaensis* (progenitor B), followed by whole-genome duplication (23–29). Two diploid progenitors diverged ∼2 million years ago (Mya), whereas divergence between each A. *hypogaea* subgenome and its corresponding progenitor occurred ∼0.2 Mya (28). This short evolutionary timespan has resulted in two subgenomes being clearly distinguishable (25), and likely insufficient time for extensive *cis*-regulatory divergence between derived and ancestral states. Peanut, therefore, represents an exceptional model to examine the early stages of *cis*-regulatory evolution and to trace how regulatory elements evolve through genome hybridization and duplication.

To investigate sequence variation in *cis*-regulatory elements and its relationship to regulatory activity and gene expression divergence, we analyzed three peanut species: *A. hypogaea, A. duranensis* and *A. ipaensis*, using both leaf and root tissues. We integrated Assay for Transposase Accessible Chromatin Sequencing (ATAC-seq) to map genome-wide chromatin accessibility, Chromatin Immunoprecipitation Sequencing (ChIP-seq) to profile the histone modifications, and RNA-seq to quantify transcript abundance. We identified Accessible Chromatin Regions (ACRs) and classified them into distinct evolutionary categories based on sequence conservation. Comparative analyses of sequence variation, chromatin accessibility states, histone modification patterns and CNSs dynamics, revealed distinct evolutionary and functional signatures among these classes. Together, these results illustrate the dynamic regulatory landscapes of recently formed polyploids, where ACRs from different evolutionary origins follow diverse trajectories that collectively shape regulatory balance and expression diversification within the polyploid genome.

## Results

### Mapping the accessible chromatin landscape in peanut

We generated 12 ATAC-seq libraries, comprising two biological replicates of leaf and root tissues from each of the three peanut species: *A. hypogaea, A. duranensis* and *A. ipaensis*. An average of 84,513,689 raw reads were obtained per library. After adapter trimming and quality filtering, uniquely mapped reads were used for peak calling, and regions meeting defined thresholds were designated as ACRs (see SI Methods). On average 61,539 peaks were identified per replicate (**Table S1**). The average proportion of reads around Transcription Start Site (TSS) and average Fraction of Reads in Peak (FRiP) score were 0.35 and 0.48, indicating the good quality of ATAC-seq data (**Table S1**).

### Comparative characterization of ACRs across four peanut genomes

We merged ACRs identified from leaf and root tissues for each genome: subgenome A and subgenome B of *A. hypogaea* and the two diploid progenitors, *A. duranensis* and *A. ipaensis*, yielding 56,526, 56,847, 113,025, and 95,584 ACRs, respectively (**Table S2**). These merged ACRs were used as references for differential chromatin accessibility analysis between leaf and root tissues. Regions showing no significant difference in chromatin accessibility were classified as broad ACRs, whereas those with significantly higher chromatin accessibility in either tissue were designated as leaf-specific or root-specific (**Fig. 1A**). These tissue-specific labels (broad, and leaf-specific or root-specific) were retained and used throughout downstream analyses.

**Figure 1.**
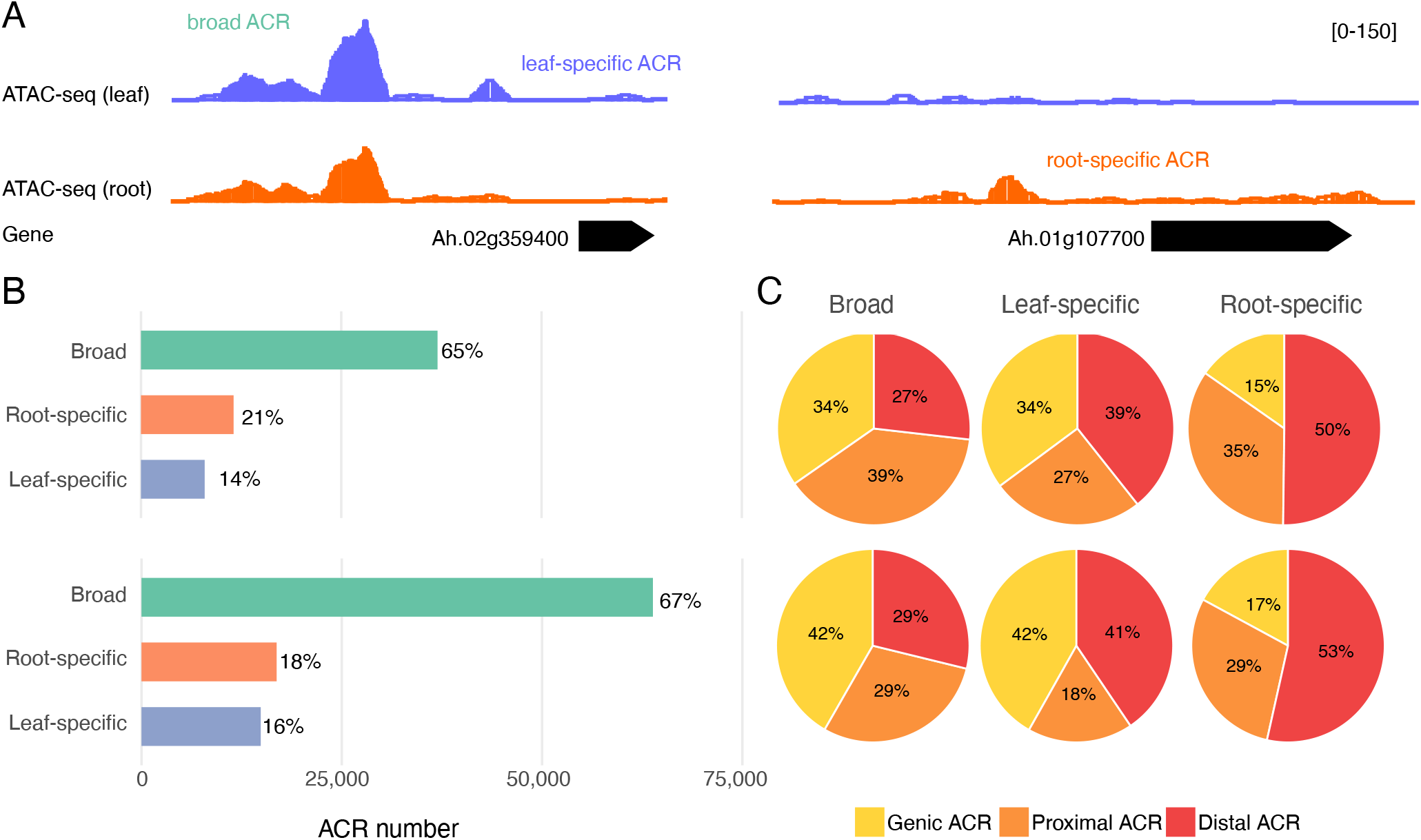
Characterization of ACRs identified in peanut subgenome and diploid progenitor. **(A)** Screenshot illustrating example of tissue-specific ACRs and broad ACRs. **(B)** Number and proportion of tissue-specific versus broad ACRs in subgenome B of *A. hypogaea* (top) and progenitor B (*A. ipaensis*) (bottom). **(C)** Distribution of genic, proximal and distal ACRs within broad, leaf-specific and root-specific categories in subgenome B (top) and progenitor genome B (bottom).

Broad ACRs accounted for the majority of ACRs, accounting for 65.4%-72.8%, with tissue-specific ACRs representing a smaller fraction (11.4%-20.5%) (**Fig. 1B, Fig. S1** and **Table S2**), consistent with findings from previous studies (13, 14, 30–32). Based on proximity to genes, ACRs were further categorized as genic, proximal (≤2 kb from genes), or distal (>2 kb). Among tissue-specific ACRs, distal regions were most prevalent (37.1%-55.8%), whereas broad ACRs were enriched in genic and proximal regions, together accounting for more than 70% (**Fig. 1C, Fig. S1** and **Table S2**). Although a greater number of ACRs were identified in diploid genomes, both diploids and two subgenomes exhibited comparable ACR patterns of tissue specificity and genomic distribution (**Fig. 1B-1C, Fig. S1** and **Table S2**).

### Identification of ACRs with distinct evolutionary trajectories

The well-resolved phylogenetic relationships between the two subgenomes and their respective diploid progenitors provide a rigorous framework for tracing the evolutionary trajectories of *cis*-regulatory elements (**Fig. 2A**). To infer the sequence ancestry of ACRs in polyploid peanut, we performed BLAST searches of ACRs from each subgenome against the other subgenome and both diploid progenitor genomes. To preserve the local genomic context of each regulatory region, we implemented a synteny-based BLAST pipeline adapted from (13). In this framework, each query ACR was anchored by its flanking homoeologous genes, and BLAST searches were restricted to the corresponding syntenic intervals in the target genomes (**Fig. S2**). We focused on ACRs located within syntenic intervals conserved across all four genomes: the two subgenomes and two diploid genomes, resulting in 33,751 and 30,994 ACRs (approximately 60% of the total identified ACRs) from subgenome A and subgenome B, respectively, being BLASTed. Although restricting analyses to conserved syntenic intervals may exclude some non-collinear homologous ACRs and transposable elements (TEs)-mediated relocations (33, 34), it provides a robust baseline for tracking regulatory evolution. Based on sequence conservation profiles, ACRs were classified into five evolutionary categories reflecting their possible origins: 1) **subgenome-specific**, present only in one subgenome and absent from the other subgenome and both diploid genomes, suggesting potential *de novo* emergence; 2) **polyploid-specific**, shared between subgenomes but not detected in diploids, indicating formation after hybridization but prior to genome duplication or less likely, simultaneous loss in both progenitors; 3) **lineage-specific**, conserved only between subgenome and its corresponding diploid progenitor, representing inheritance from the ancestral genome; 4) **peanut-conserved**, retained across four genomes, indicating long-term retention throughout the peanut lineage; 5) **potential peanut-conserved**, detected in three genomes or shared between a subgenome and the non-corresponding diploid genome (**Fig. 2A**).

**Figure 2.**
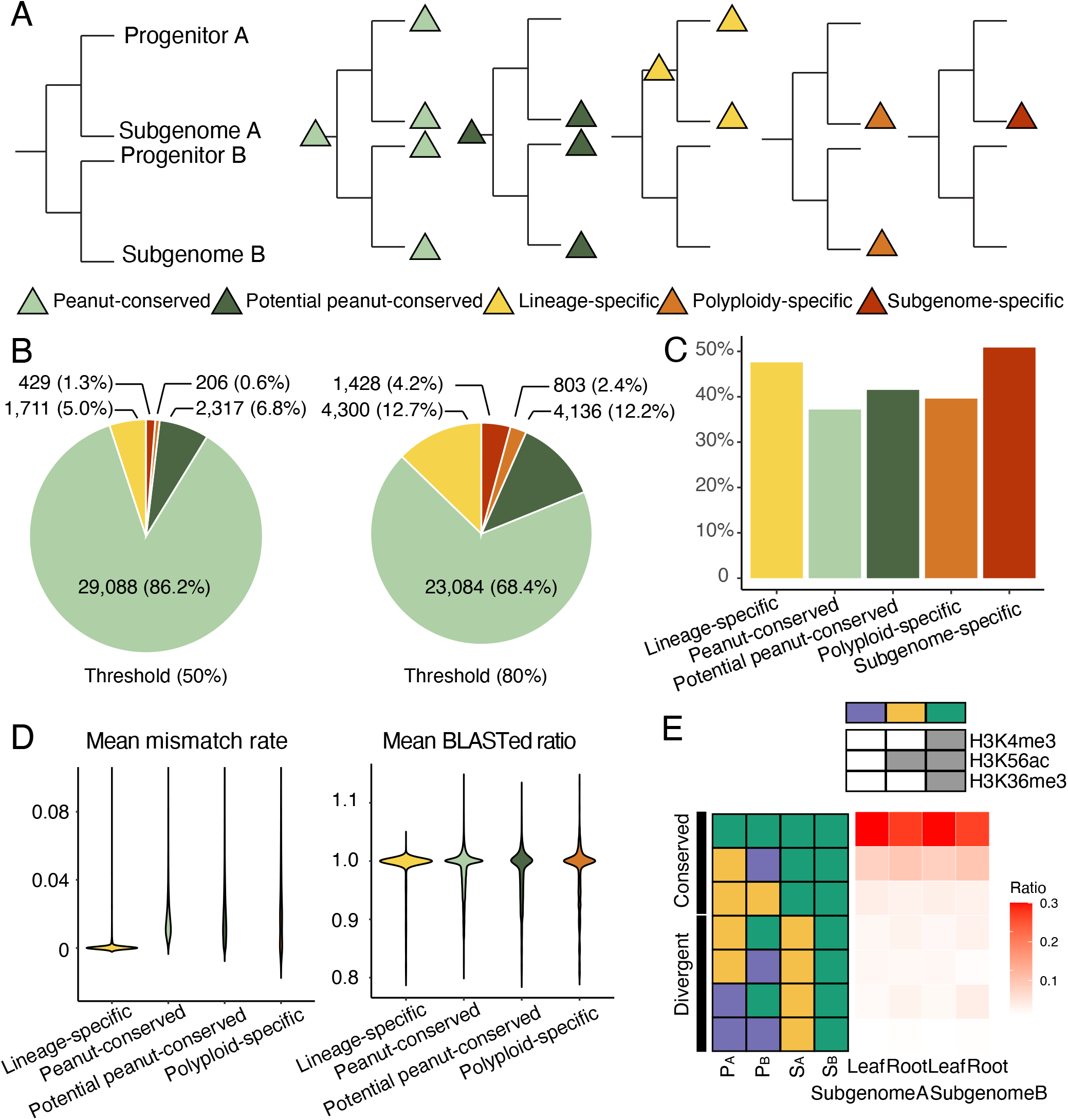
Identification and characterization of ACRs with distinct evolutionary trajectories. **(A)** Schematic illustrating the classification of ACRs based on sequence conservation profiles. Peanut-conserved ACRs are retained across all four genomes, the two subgenomes of polyploid peanut and both diploid progenitors, reflecting ancestral conservation predating progenitor divergence; potential peanut-conserved ACRs are detected in three genomes, or between a subgenome and a non-corresponding progenitor; lineage-specific ACRs are shared between a subgenome and its corresponding progenitor, representing inheritance from the respective diploid lineage; polyploid-specific ACRs likely originated after hybridization but before whole-genome duplication or less likely, simultaneous loss in both progenitors; subgenome-specific ACRs are unique to one subgenome, suggesting potentially *de novo* emergence following polyploidization. **(B)** Distribution of ACRs with distinct evolutionary trajectories (identified in subgenome A) under two sequence-similarity thresholds, using BLASTed ratios ≥50% (left) and ≥80% (right) to define sequence conservation. **(C)** Proportion of each evolutionary ACR category overlapping with TEs (in subgenome B), with the criteria of at least 20 bp overlap. **(D)** Genetic variation across peanut-conserved, potential peanut-conserved, lineage-specific, and polyploid-specific ACRs (in subgenome B), measured by mean mismatch rate and mean BLASTed ratio. **(E)** Distribution of distinct histone modification patterns in conserved ACRs and their corresponding BLASTed regions across all four genomes. The top panel summarizes the three histone modification states (purple, blue and green bars) defined by combinations of the histone modification marks, H3K4me3, H3K56ac and H3K36me3, with grey bars indicating the presence of each respective mark. The bottom-left schematic illustrates the classification of histone modification patterns shown from top to bottom: (i) patterns conserved across all four genomes; (ii) patterns shared between the two subgenomes but different in both progenitors; (iii) patterns shared between both subgenomes and both progenitors but differing between subgenome and progenitor groups; (iv) patterns distinct between the two subgenomes but each matching its corresponding progenitor; (v) patterns consistent only between subgenome A and progenitor A; (vi) patterns consistent only between subgenome B and progenitor B; and (vii) patterns different from both progenitors. The top three categories represent conserved histone modification patterns between the two subgenomes, whereas the remaining four represent divergent patterns. The bottom-right heatmap shows the proportion of each histone modification pattern in leaf and root tissues within subgenome A and subgenome B. P_A_: progenitor A; P_B_: progenitor B; S_A_: subgenome A and S_B_: subgenome B.

To evaluate the stability of these classifications, we applied two sequence similarity thresholds (≥50% and ≥80%), requiring BLAST alignments of at least 75bp or 120bp, respectively. Raising the threshold from 50% to 80% led to a marked reclassification of ACRs, particularly with many shifting from peanut-conserved to lineage-specific categories (**Fig. 2B, Fig. S3** and **Table S3**). For example, in subgenome A, the proportion of lineage-specific ACRs increased from 5.0% to 12.7% whereas the proportion of peanut-conserved ACRs decreased from 86.1% to 68.3% (**Fig. 2B, Fig. S3** and **Table S3**). This shift indicates that sequence divergence has accumulated within regions previously considered conserved, reflecting their recent evolutionary trajectories. Because the higher threshold more accurately captures these recent differences, subsequent analyses were based on ACR classifications defined by the 80% criterion. In both subgenomes, most ACRs within conserved syntenic intervals were peanut-conserved (∼68%) (**Fig. 2B, Fig. S3** and **Table S3**). When including potential peanut-conserved ACRs, this proportion increased to approximately 80%, underscoring their overall high sequence conservation. Lineage-specific accounted for ∼12%, followed by subgenome-specific (∼2-4%) and polyploid-specific ACRs (∼2-4%) (**Fig. 2B, Fig. S3** and **Table S3**). Notably, subgenome-specific ACRs were more abundant in subgenome A (4.2%) than in subgenome B (2.6%) (**Fig. S3** and **Table S3**).

Tissue specificity patterns were largely consistent across categories, with most ACRs (∼68%) exhibiting broad chromatin accessibility (**Fig. S4** and **Table S4**). Genomic distributions were also comparable across categories, where the largest proportions were proximal ACRs (∼38%), followed by distal ACRs (∼30%) and then genic ACRs (∼24%), although peanut-conserved ACRs, potential peanut-conserved ACRs and polyploid-specific ACRs had slightly higher proportions of genic ACRs (∼35%) (**Fig. S4** and **Table S4**).

### TEs disproportionately shape ACRs with different evolutionary trajectories

Repeat sequences are abundant in the peanut genome, with retroelements and DNA transposons comprising the majority (29). TEs are well known to drive the emergence of novel *cis*-regulatory elements by introducing regulatory sequences into new genomic contexts (5, 12, 33–37). In our analysis, TE associations were more frequent among subgenome-specific and lineage-specific ACRs (**Fig. 2C, Fig. S5** and **Table S4**). For instance, 50.9% of subgenome-specific ACRs in subgenome B overlapped with repeat elements by at least 20 bp (TE-mediated ACRs), and 17.6 % were entirely embedded within ACRs (**Fig. 2C, Fig. S5** and **Table S4**), underscoring a strong contribution of TEs to regulatory novelty. Both TE-mediated ACRs and ACRs containing fully embedded TEs were more abundant in subgenome B than in subgenome A across most ACR categories (**Fig. S5-S6** and **Table S4**). Although subgenome-specific ACRs represented an exception in terms of absolute counts, the proportion of TE-mediated or TE fully embedded ACRs relative to all subgenome-specific ACRs remained higher in subgenome B than in subgenome A (**Fig. S5-S6** and **Table S4**).

Among TE-mediated ACRs, the four most abundant TE families were consistently Retro, Mu, CACTA and LINE, in descending order, across ACRs with different evolutionary trajectories in both subgenomes (**Fig. S7** and **Table S5**). In contrast, TE fully embedded ACRs exhibited different patterns between the two subgenomes. In subgenome A, Feral contributed the second-largest proportion, whereas LINE was the second most abundant family in subgenome B, with Retro elements remaining the dominant contributors in both subgenomes (**Fig. S7** and **Table S5**).

We further compared the chromatin accessibility between the TE-mediated and non-TE-mediated ACRs across both subgenomes. TE-mediated ACRs exhibited significantly higher chromatin accessibility than non-TE-mediated ACRs (adjusted p-values ≈ 0), whereas differences in chromatin accessibility between TE-mediated ACRs from subgenome A and subgenome B were not significant or only marginally significant (adjusted p-values >0.5 or ≈ 0.4) (**Fig. S8** and **Table S6**).

Collectively, these results indicated that TEs contribute to the evolution of regulatory regions and are associated with elevated chromatin accessibility. Moreover, differences in the abundance of TE-mediated ACRs and in TE family composition suggest that TE insertions may have exerted asymmetric influences on the evolution of regulatory landscapes between the two subgenomes.

### Genetic variation patterns among ACRs with distinct evolutionary histories

To evaluate sequence variation among ACRs with different evolutionary trajectories, we quantified two metrics: the BLASTed ratio, defined as the length of the query ACR divided by the length of its BLASTed sequence in the corresponding reference region, and the mismatch rate, defined as the number of mismatches per BLASTed sequence. Subgenome-specific ACRs were excluded from this analysis because no corresponding sequences exist in the reference regions. Across the remaining categories, most ACRs exhibited high BLASTed ratios (median ∼0.98 ) and low mismatch rates (median < 0.02) (**Fig. 2D, Fig. S9** and **Table S4**). These values contrast sharply with those reported for conserved ACRs in soybean, where the median BLASTed ratio of recently duplicated ACRs was ∼0.75 and the median mismatch rates was ∼0.05 (22). This comparison underscores the elevated regulatory sequence conservation in this recently formed polyploids and highlights the influence of divergence time on regulatory sequence decay.

Interestingly, lineage-specific ACRs exhibited the highest BLASTed ratios and lowest mismatch rates, whereas polyploid-specific ACRs displayed the lowest BLASTed ratios and highest mismatch rates, with peanut-conserved and potential peanut-conserved falling in between (**Fig. 2D, Fig. S9** and **Table S4**). Across these comparisons, lineage-specific ACRs were consistently and significantly different from the other groups (**Table S7**). Compared with peanut-conserved and potential peanut-conserved ACRs, the lower sequence divergence of lineage-specific ACRs likely reflects their more recent emergence. However, it is somewhat counterintuitive that lineage-specific ACRs were less divergent than polyploid-specific ACRs, as lineage-specific ones may have arisen earlier and might be expected to accumulate more mutations over time. The elevated conservation of lineage-specific ACRs may instead reflect stronger selective pressure associated with their single-copy status in the polyploid genome. In contrast, the duplicated nature of polyploid-specific ACRs could relax selective constraints, facilitating greater sequence divergence, although we cannot completely rule out the possibilities that some of these lineage-specific ACRs originated early but were lost simultaneously in both diploid progenitors, a scenario that is likely rare.

### Histone modification dynamics of sequence-conserved ACRs

To characterize the histone modification landscape associated with ACRs, we performed ChIP-seq for three peanut species in both leaf and root tissues. Chromatin state conservation was inferred from three active histone marks: H3K4me3, H3K56ac, and H3K36me3. We did not profile repressive histone marks or DNA methylation, which may exhibit different regulatory dynamics (38, 39). Genome-wide patterns of histone marks were consistent with previous reports (12, 40– 43), with H3K4me3 and H3K56ac enriched around transcription start sites, and H3K36me3 enriched within gene bodies (**Fig. S10**). We then intersected ACR coordinates with peaks of individual histone marks and classified ACRs into three states: transcribed (signals present across all three marks, indicative of active transcription), Kac-enriched (signal exclusively for H3K56ac, potentially reflecting enhancer capacity), and unmodified (no signal for any of the examined histone marks) (**Fig. 2E**), following the clustering criteria described in (12).

Comparison of histone states across peanut-conserved ACRs and their corresponding BLASTed regions revealed that most ACRs (∼85%) retained identical histone states between the two subgenomes, and the majority also shared the same states as both diploid progenitors (∼60%) (**Fig. 2E** and **Table S8**). A small subset (∼6%) displayed conserved states within subgenomes and within progenitors but differed between subgenomes and progenitors (**Fig. 2E** and **Table S8**). The remaining ∼15% exhibited divergent histone states between subgenomes; in most of these cases, one or both subgenomes matched the histone state of the corresponding progenitor, highlighting strong parental-legacy effects, whereas only ∼2% showed entirely novel histone states absent from either progenitor (**Fig. 2E** and **Table S8**). Lineage-specific and polyploid-specific ACRs also displayed high histone state similarity, ranging from 70% to 86%, either between a subgenome and its corresponding progenitor or between the two subgenomes (**Table S8**).

### Varied chromatin accessibility bias among sequence-conserved ACRs

To further investigate the regulatory dynamics of sequence-conserved ACRs, we quantified chromatin accessibility by counting reads overlapping each conserved ACR and its corresponding BLASTed sequences. Peanut-conserved ACRs, those with highly conserved sequences across all four genomes, provide a framework for examining chromatin accessibility dynamics in both ancestral and derived states. Principal component analysis (PCA) of the normalized chromatin accessibility counts revealed a strong parental-legacy effect on chromatin accessibility variation. Principal Component 1 (PC1), explaining ∼36.3% of the total variance, separated the two lineages, subgenome A clustered with *A. duranensis*, and subgenome B with *A. ipaensis*. PC2, accounting for ∼27.5% of the variance, distinguished tissue types (**Fig. 3A** and **Fig. S11**). Following the conceptual framework of homoeolog expression bias, in which homoeologous genes exhibit differential expression (44–48), we examined chromatin accessibility bias in peanut-conserved ACRs to determine whether chromatin accessibility differed between subgenomes, whether the bias favored subgenome A or subgenome B, and whether such bias reflected parental legacy or regulatory novelties that emerged following genome hybridization and duplication. Accordingly, each ACR was classified into three major groups: (i) **parental condition**, no-bias across all four genomes or a bias consistent with the progenitors; (ii) **no bias in progeny**, a bias present in progenitors but absent in the polyploid; and (iii) **novel bias in progeny**, bias distinct from the parental condition (**Fig. 3B**). Within each category, ACRs were further designated as subgenome A- or subgenome B-biased (or progenitor A- or progenitor B-biased) (**Fig. 3B**), depending on which genome exhibited significantly higher chromatin accessibility (See Methods).

**Figure 3.**
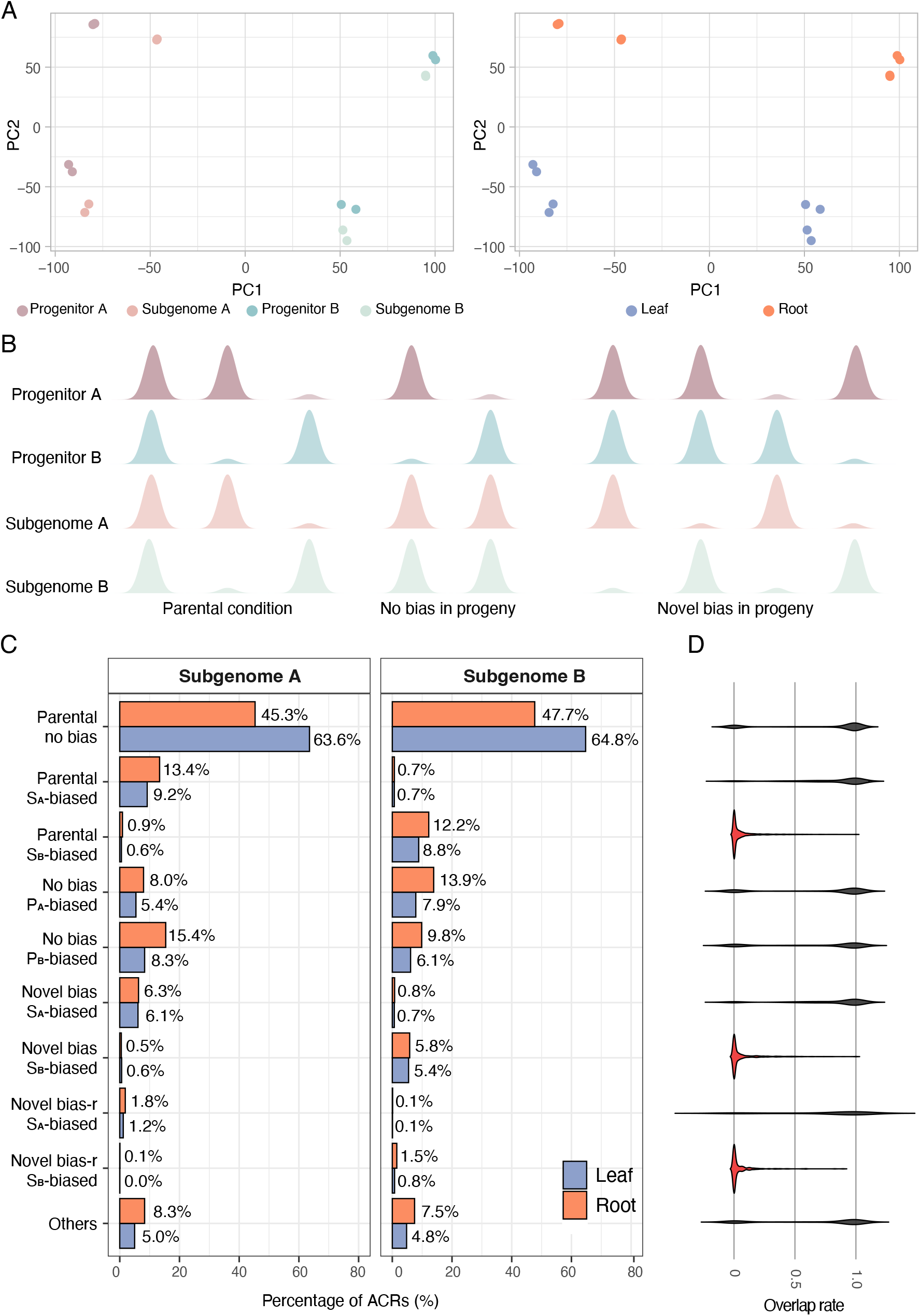
Characterization of chromatin accessibility bias among peanut-conserved ACRs. **(A)** Principal component analyses (PCA) of normalized chromatin accessibility counts for peanut-conserved ACRs (in subgenome A) across all four genomes, the two subgenomes of polyploid peanut and both diploid progenitors (left), and across two tissue types, leaf and root (right). **(B)** Schematic illustrating classification of chromatin accessibility bias based on the relative chromatin accessibility patterns between subgenomes and between their corresponding progenitors. Categories include (from left to right): parental no bias (no bias between either subgenomes or progenitors); parental subgenome A- or subgenome B-biased (bias direction in subgenome consistent with that in progenitors); no bias progenitor A- or progenitor B-biased (no bias between subgenomes but bias present between progenitors); novel bias subgenome A- or subgenome B-biased (bias emerging between subgenomes despite no bias between progenitors); and novel bias-reverse (novel bias -r) subgenome A- or subgenome B-biased (bias direction opposite the observed between progenitors). **(C)** Distribution of these chromatin accessibility bias patterns for peanut-conserved ACRs identified in subgenome A and subgenome B across leaf and root tissues. **(D)** Overlap rate between BLASTed regions of peanut-conserved ACRs (identified in subgenome B) in subgenome A and the ACRs experimentally identified in subgenome A; red labels denote subgenome B-biased ACRs. P_A_: progenitor A; P_B_: progenitor B; S_A_: subgenome A and S_B_: subgenome B.

Distributions of ACRs with different chromatin accessibility bias patterns were largely similar between leaf and root tissues, except that a greater proportion exhibited a parental no bias pattern in root tissue (**Fig. 3C** and **Table S9**). Across tissues, the majority of ACRs (∼70–80%) showed no chromatin accessibility bias between the two subgenomes. Within this group, most ACRs (∼45–63%; over 10,000 ACRs in root and over 13,000 ACRs in leaf) also lacked bias between the diploid progenitors (parental no bias), whereas the remainder (∼13–23%) exhibited progenitor-specific bias (no bias progenitor A- or progenitor B-biased) (**Fig. 3C** and **Table S9**). Approximately 20% of ACRs exhibited significant accessibility bias between subgenomes, corresponding at least 3,400 ACRs. These biased ACRs primarily reflected parental-legacy effects (parental subgenome A- or subgenome B-biased ∼10%), followed by novel bias (novel bias subgenome A- or subgenome B-biased ∼6%) and novel reverse bias (novel bias-r subgenome A- or subgenome B-biased ∼2%), in which the direction of bias was opposite to that observed in the progenitors (**Fig. 3C** and **Table S9**). ACRs biased toward subgenome A were predominantly found in subgenome A (20%) and were rare in subgenome B (1%), with a corresponding pattern for subgenome B-biased ACRs. (**Fig. 3C** and **Table S9**). We also quantified log_2_ fold change to assess the magnitude of accessibility differences. Within each subgenome, both subgenome A- and subgenome B-biased ACRs exhibited similar fold-change levels across parental-, novel- or novel reverse-biased categories (**Fig. S12**). However, ACRs biased toward their resident subgenome consistently showed higher absolute values than ACRs biased toward the opposite subgenome. For example, in subgenome A, subgenome A-biased ACRs had higher absolute fold-change values than subgenome B-biased ACRs (**Fig. S12**).

To characterize these subgenome-biased ACRs further, we overlapped the BLASTed coordinates of ACRs identified from one subgenome (e.g., subgenome B) in the other subgenome (e.g., subgenome A) with the coordinates of experimentally identified ACRs in the other subgenome (e.g., subgenome A). ACRs biased toward their resident subgenome exhibited markedly lower overlap, whereas ACRs with other chromatin accessibility bias patterns showed much higher overlap (**Fig. 3D, Fig. S13**, and **Table S10**). Normalized chromatin accessibility counts for these subgenome-biased ACRs were also substantially higher in their native subgenome compared to their BLASTed counterparts in the other subgenome (**Table S11**). Together, these results indicate that subgenome-biased ACRs are largely restricted to their respective subgenome, but typically not detected as ACRs in the other subgenome, suggesting differential chromatin accessibility between subgenomes often manifests as presence-absence variation rather than as quantitative shifts.

We next examined sequence variation among peanut-conserved ACRs exhibiting different chromatin accessibility bias patterns. Although no strong overall pattern was observed, parental no bias ACRs consistently had the lowest mismatch rates and relatively high BLASTed ratio compared to ACRs with altered chromatin accessibility (**Table S12**), suggesting that sequence divergence contributes to chromatin accessibility variation.

Comparisons of lineage-specific ACRs between each subgenome and its corresponding diploid progenitor, as well as of polyploid-specific ACRs between subgenomes, also revealed that most did not exhibit significant differences in chromatin accessibility (**Table S13**), consistent with previous observations of histone states (**Fig. 2E** and **Table S8**).

### Age-dependent conservation and loss of CNSs in peanut subgenomes and progenitors

*Cis*-regulatory elements are often highly conserved, reflecting selection to maintain essential regulatory function (10, 49–53). To gain further insights into their evolutionary dynamics, we examined CNSs in the context of polyploid peanut and its corresponding diploid progenitors, providing complementary perspective to studies that have largely overlooked lineage-specific regulatory evolution (10, 36, 54, 55). CNS dataset used in this study was obtained from Conservatory (www.conservatorycns.com) and was generated using a gene-centric CNS discovery algorism that accounts for complex orthology relationships, structural variation and rapid sequence divergence (10).

Consistent with previous findings (10), ancient CNSs, those conserved at or above the Angiosperm level, were less numerous but exhibited higher copy numbers and greater retention than lineage CNSs, such as family-level CNSs. For example, 71% of CNSs were conserved at the Fabaceae level, whereas only ∼2% were conserved at the Angiosperm level (**Fig. 4A** and **Table S14**); ∼75% of Tracheophyte-level (vascular plants) CNSs had at least two copies, compared with 25% of Fabaceae-level CNSs (**Fig. 4B** and **Table S15**); and 55% of Tracheophyte-level CNSs were retained across all three peanut species, versus 23.9% of Fabaceae-level CNSs (**Fig. 4C** and **Table S16**). In contrast to the previously proposed view that one paralog largely retains ancestral CNSs while the other undergoes rapid turnover (10), we found that ∼42% of Tracheophyte- and Angiosperm-level CNSs were preserved across both subgenomes and their diploid progenitors (**Fig. 4C** and **Table S16**). This pattern may partly reflect the relatively recent origin of the allotetraploid, leaving insufficient time for ancient CNSs to diverge.

**Figure 4.**
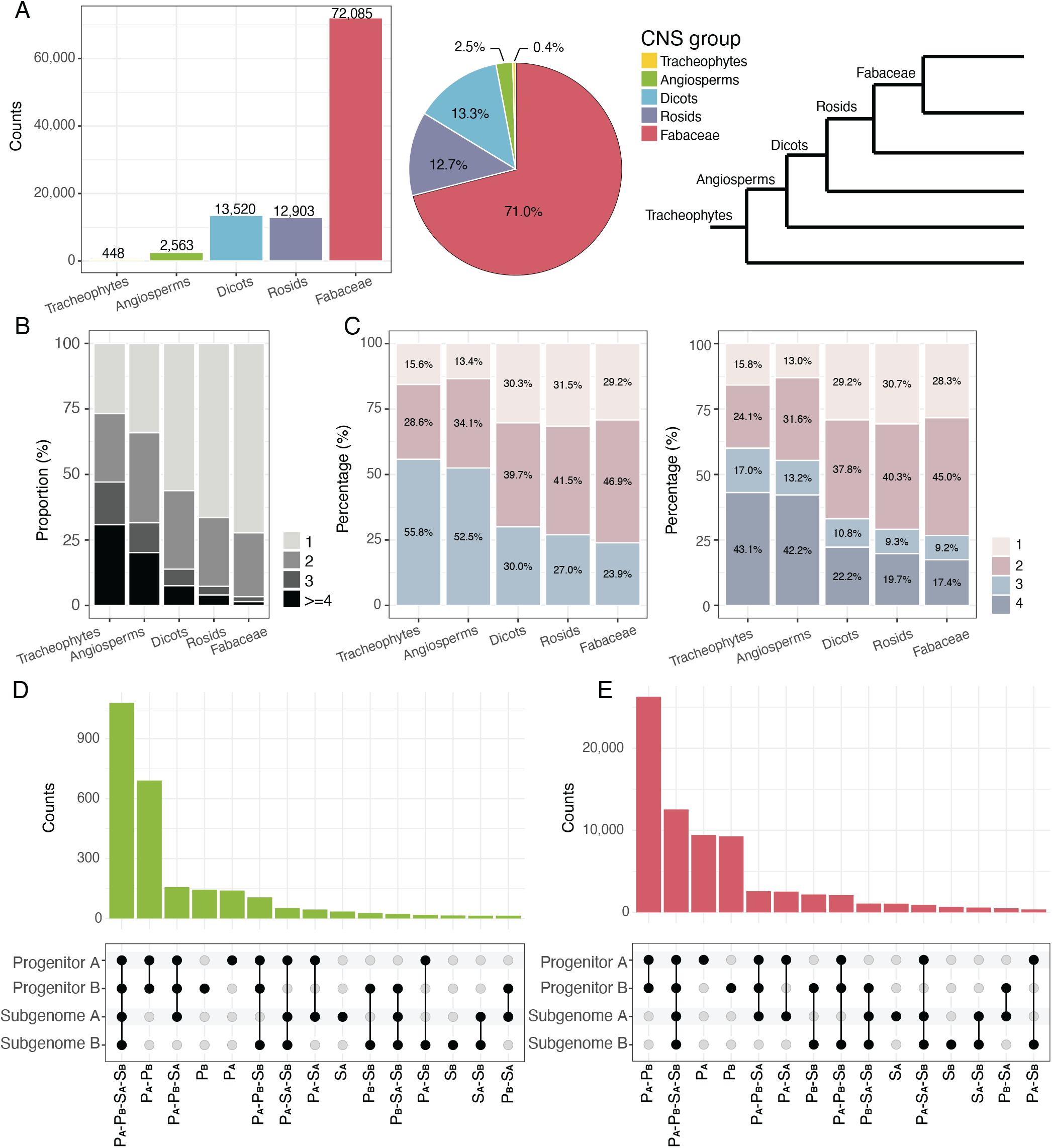
Characterization of CNS dynamics within the polyploid-diploid peanut framework. **(A)** Number and proportion of CNSs identified in the three peanut species, grouped by their evolutionary age of conservation, from ancient Tracheophyte-level or Angiosperm-level CNSs to increasingly lineage-restricted CNSs in Dicots, Rosids, and Fabaceae. **(B)** Distribution of CNS copy numbers across CNSs with different evolutionary age of conservation. **(C)** Distribution of CNSs with different conservation levels across the three peanut species (left) and across the four genomes, the two subgenomes of polyploid peanut and the two diploid progenitors (right), for each CNS evolutionary age of conservation category. Numbers from 1 to 4 indicate CNSs conserved in one, two, three, or all four species/genomes. **(D-E)** Complex spectrum of CNS dynamics (CNS conservation states) across the diploid-polyploid framework for Angiosperm-level CNSs (D) and Fabaceae-level CNSs (E). CNSs are categorized as: (i) conserved across all four genomes, (ii) conserved across three genomes, (iii) conserved exclusively in progenitors, (iv) conserved exclusively in polyploid subgenomes, (v) conserved between a subgenome and its corresponding progenitor, (vi) conserved between a subgenome and the non-corresponding progenitor, and (vii) restricted to a single genome. P_A_: progenitor A; P_B_: progenitor B; S_A_: subgenome A and S_B_: subgenome B.

Beyond these general trends, we observed a more complex spectrum of CNS dynamics within the diploid-polyploid framework. Based on their presence across the four genomes, CNSs were classified into seven categories : (i) conserved across all four genomes, (ii) conserved across three genomes, (iii) conserved exclusively in progenitors, (iv) conserved exclusively in polyploid subgenomes, (v) conserved between a subgenome and its corresponding progenitor, (vi) conserved between a subgenome and the non-corresponding progenitor, and (vii) restricted to a single genome (**Fig. 4D-4E** and **Fig. S14**). Categories conserved in only a subset of genomes may reflect loss from the remaining genomes.

We then examined whether the distributions of these CNS conservation states varied with CNS age. Ancient CNSs were dominated by those conserved across all four genomes, followed by those retained only in progenitors (**Fig. 4D** and **Fig. S14**). In contrast, lineage CNSs were most frequently conserved exclusively in progenitors, with the second most category being those conserved across all four genomes (**Fig. 4E** and **Fig. S14**). The next predominant groups within lineage CNSs included CNSs restricted to progenitor A or progenitor B, as well as CNSs absent in one subgenome but present in the other three genomes (**Fig. 4E** and **Fig. S14**). These categories were also among the next common for ancient CNS (**Fig. 4D** and **Fig. S14**); however, progenitor-restricted CNSs were substantially more frequent than subgenome-missing CNSs among lineage CNSs (**Fig. 4D** and **Fig. S14**), whereas this disparity was minimal for ancient CNSs (**Fig. 4E** and **Fig. S14**). These patterns indicate that CNSs of different evolutionary ages follow distinct trajectories: ancient CNSs tend to be stably retained, whereas lineage CNSs exhibit substantial turnover.

On the other hand, the large number of lineage CNSs present in one or both progenitors but absent from the polyploid (**Fig. 4E** and **Fig. S14**) suggests widespread loss of these elements after hybridization and polyploidization. This may reflect the inherent dynamics of *cis*-regulatory elements: some regulatory elements, possibly lineage CNSs, that are initially conserved in both progenitors and the polyploid, may rapidly evolve into new sequences and diverge enough to escape detection as lineage CNSs in the polyploid. Alternatively, hybridization, whole-genome duplication, and subsequent regulatory reorganization may drive the extensive loss of lineage CNSs in the polyploid. Further work will be needed to disentangle these possibilities. We also observed many CNSs conserved exclusively between each subgenome and its corresponding progenitors, particularly for lineage CNSs (**Fig. 4D-4E** and **Fig. S14**), underscoring the strong influence of parental legacy on regulatory architecture. Other categories occurred at much lower frequencies (**Fig. 4D-4E** and **Fig. S14**).

### Impacts of the CNS dynamics on chromatin accessibility variation of sequence-conserved ACRs

To explore whether CNS turnover correlates with the chromatin accessibility variation observed in sequence-conserved ACRs, we examined CNSs dynamics within these regions using a comparative framework. We applied the same presence/absence–based CNS conservation state classification described above (**Fig. 4D-4E** and **Fig. S14**), but in a more targeted context: each CNS was categorized based on its association with genes in the same orthogroup across the four genomes and its location within a sequence-conserved ACR and/or its corresponding BLASTed regions (see Methods) (**Table S17**). We then summarized the distribution of CNS conservation state for each ACR using a rank-normalized approach that accounts for both CNS conservation state and its abundance (see Methods).

We grouped ACRs by their chromatin accessibility bias patterns and examined how CNS conservation state distributions corresponded to these distinct ACR groups (**Fig. 5** and **Fig. S15**). The patterns were consistent between leaf and root tissues (**Fig. 5** and **Fig. S15**). ACRs with parental subgenome-biased chromatin accessibility were frequently associated with CNSs conserved exclusively between that subgenome and its corresponding diploid progenitor (e.g., P_B_-S_B_ in **Fig. 5**). ACRs with parental no bias chromatin accessibility tended to contain CNSs conserved across all four genomes (e.g., P_A_-P_B_-S_A_-S_B_ in **Fig. 5**). ACRs exhibiting novel bias subgenome-biased chromatin accessibility were more likely to harbor CNSs unique to that subgenome (e.g., S_B_ in **Fig. 5**) or CNSs missing in the other subgenome (e.g., P_A_-P_B_-S_B_ in **Fig. 5**). In contrast, CNSs conserved across three genomes were often not enriched in any chromatin accessibility category (e.g., P_B_-S_A_-S_B_, P_A_-S_A_-S_B_, P_A_-P_B_-S_A_ in **Fig. 5**). CNSs conserved exclusively in progenitor A (e.g., P_A_ in **Fig. 5**) or progenitor B (e.g., P_B_ in **Fig. 5**) were unexpectedly common across nearly all accessibility groups, despite they being lost in the subgenomes. CNS exclusively conserved in both progenitors (e.g., P_A_-P_B_ in **Fig. 5**) showed a distinctive pattern: they were broadly enriched but specifically depleted in parental subgenome-biased chromatin accessibility, suggesting that these ACRs carry regulatory features that differ markedly between the two progenitor lineages. Together, these results suggest that CNSs with different conservation states contribute meaningfully to chromatin accessibility bias patterns of sequence-conserved ACRs, shaping their regulatory activities.

**Figure 5.**
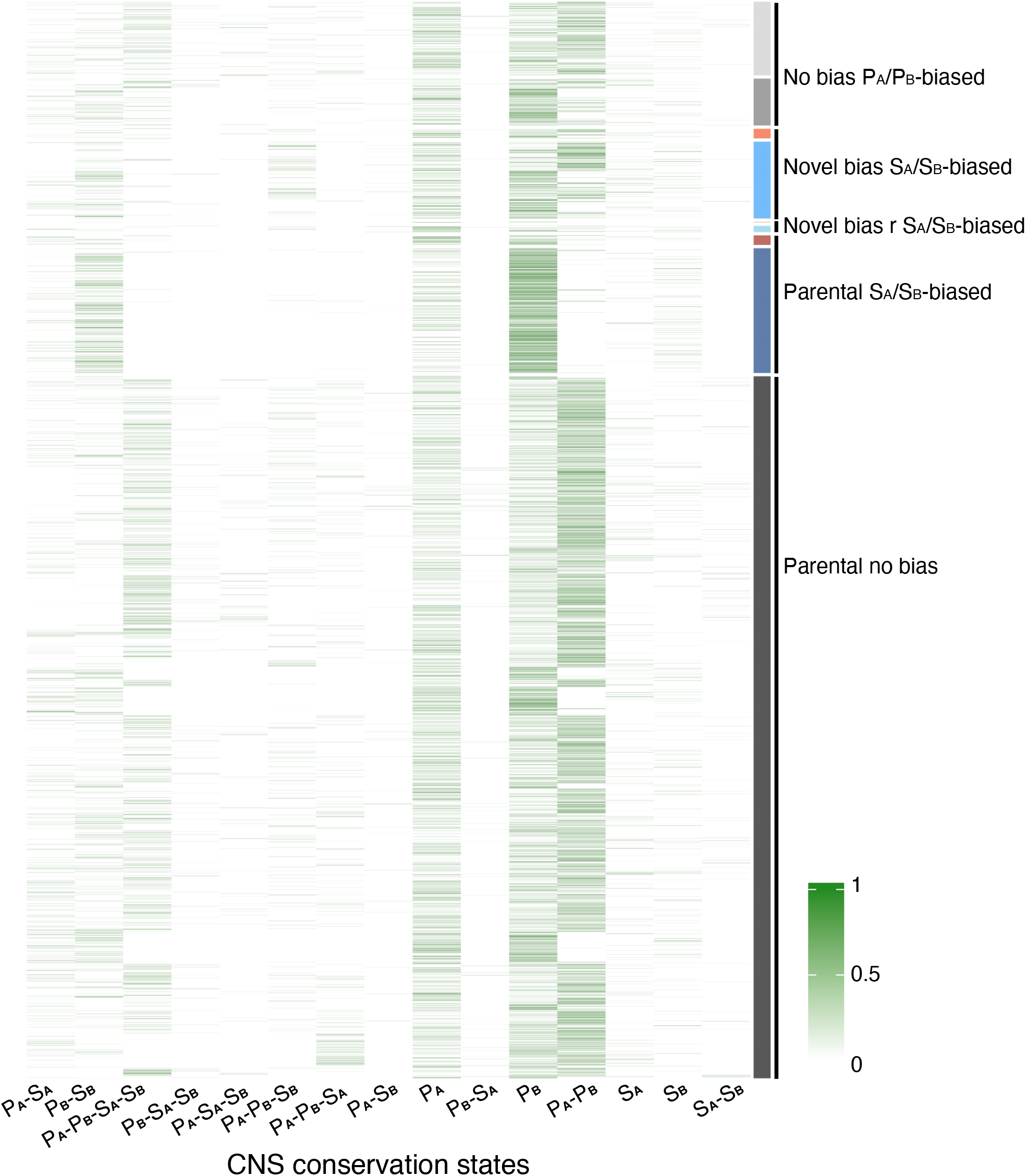
Rank-normalized scores of CNS conservation states across peanut-conserved ACRs with different chromatin accessibility bias patterns. 4E) Heatmap showing the rank-normalized scores of CNS conservation states (as defined in Fig. 4D-4E) across peanut-conserved ACRs in subgenome B, annotated according to leaf chromatin accessibility categories (as in Fig. 3C). For each ACR, CNS conservation states were summarized using a rank-normalization approach: all non-zero CNS conservation states were ranked in descending order of abundance and normalized by the number of states present, producing values between 0 and 1. A value of 1 indicates that an ACR contains CNSs exclusively from a single conservation state. Rows are grouped by ACR chromatin accessibility bias patterns (as in Fig. 3C), and the color scale reflects the relative rank-normalized score within each ACR. P_A_: progenitor A; P_B_: progenitor B; S_A_: subgenome A and S_B_: subgenome B.

We also analyzed lineage- and polyploid-specific ACRs using simplified CNS classifications. For lineage-specific ACRs, CNSs were categorized as conserved exclusively in the subgenome, in the corresponding progenitor, or in both. The distributions of the CNS with different conservation states did not differ between lineage-specific ACRs with the same versus different chromatin accessibility states (**Table S18** and **Fig. S16**). For polyploid-specific ACRs, CNSs were categorized as conserved exclusively in one subgenome or in both. Among polyploid-specific ACRs with differential accessibility, CNSs conserved in both subgenomes were consistently depleted (**Table S19** and **Fig. S17**).

### Homoeolog expression bias in peanut and its potential underlying mechanisms

To characterize homoeolog expression bias in peanut, we performed RNA-seq on leaf and root tissues from the same three peanut species, each with two biological replicates. To enable cross-genome comparisons, we focused on homoeologous genes identified across the two subgenomes and their corresponding diploid progenitors. PCA of normalized gene counts revealed that PC1 (explaining 44.3% of total variance) separated the diploid progenitors from the subgenomes, PC2 (26%) further distinguished the two subgenomes, and PC3 (19.4%) differentiated the two tissue types (**Fig. 6A**). Homoeologous genes displayed pronounced expression divergence both between subgenomes and relative to their diploid counterparts, whereas the two diploid progenitors showed highly similar expression profiles for these genes (**Fig. 6A**).

**Figure 6.**
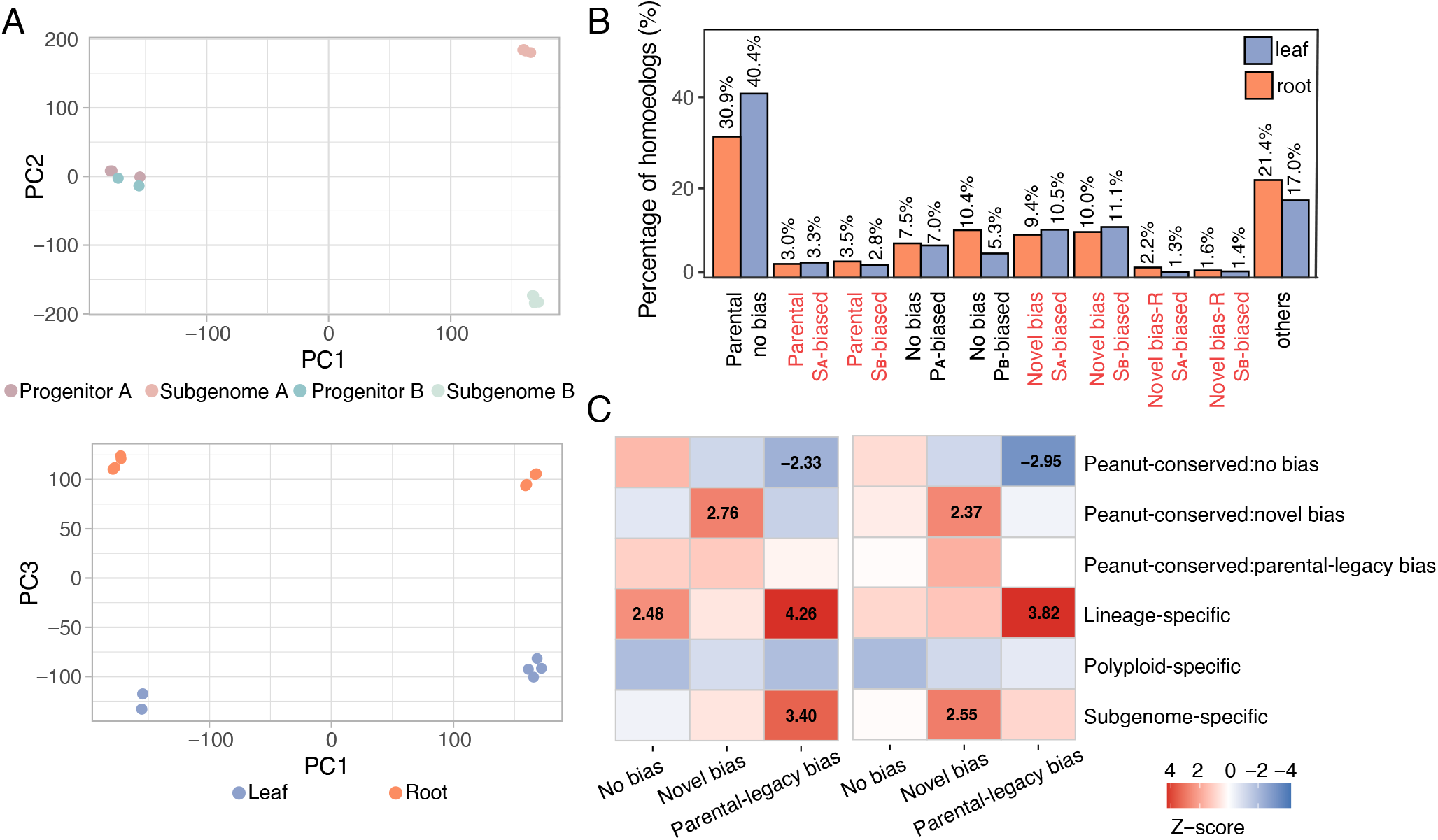
Homeolog expression bias in peanut and its association with ACRs of distinct evolutionary trajectories. **(A)** Principal component analyses (PCA) of normalized expression counts for homoeologous genes across all four genomes, the two subgenomes of polyploid peanut and both diploid (top), and across two tissue types, leaf and root (bottom). **(B)** Distribution of homeolog expression bias categories across leaf and root tissues. Biased expression categories were labelled with red font. **(C)** Heatmap showing enrichment patterns between expression types (no-bias, novel bias and parental-legacy bias) and ACR evolutionary types including peanut-conserved:no bias, peanut-conserved:novel bias, peanut-conserved:parental-legacy bias, lineage-specific, polyploid-specific and subgenome-specific ACRs. Absolute Z-scores greater than 2 are indicated.

Using the same conceptual framework applied to classify chromatin accessibility bias patterns we categorized homoeolog expression bias (**Fig. 3B**). The overall distributions of these categories were highly consistent between leaf and root tissues **(Fig. 6B** and **Table S20**). More than 50% of genes exhibited no expression bias between the two subgenomes, with the majority showing a parental no bias pattern (**Fig. 6B** and **Table S20**), consistent with previous findings (44). Among genes exhibiting bias between the two subgenomes (∼3,500 genes), novel bias accounted for the largest proportion (∼20%), followed by parental-legacy effects (∼6%), and novel bias reverse (∼3%) (**Fig. 6B** and **Table S20**). These patterns suggest that most expression asymmetries might have arisen *de novo* following hybridization and polyploidization, rather than being inherited from the ancestral diploid progenitors.

To investigate how ACRs with different evolutionary trajectories relate to homoeolog expression bias, we classified both homoeolog expression patterns and ACRs into defined categories. Homoeolog expression patterns were simplified into four classes: no-bias, novel-bias, parental-legacy bias and others (**Fig. 6C**). ACRs were first grouped by sequence conservation as peanut-conserved, potential peanut-conserved, lineage-specific, polyploid-specific and subgenome-specific and peanut-conserved ACRs were further subdivided by chromatin accessibility bias into no-bias (peanut-conserved:no bias), novel-bias (peanut-conserved:novel bias), parental-legacy bias (peanut-conserved:parental-legacy bias), and others (peanut-conserved:others) (**Fig. 6C**). To ensure robust evolutionary inference, we restricted the analysis to homoeologous genes associated with ACRs that could be confidently classified based on BLAST results, excluding those had not been BLASTed due to their absence from conserved syntenic regions. This integrative framework allowed us to systematically examine whether specific classes of regulatory regions are preferentially associated with distinct modes of homoeolog expression divergence.

We quantified the number of ACRs within each category for every orthogroup. When comparing these counts across orthogroups with different homoeolog expression bias patterns, only peanut-conserved:no bias consistently exhibited an average of more than one ACR per orthogroup (**Fig. S18**). Lineage-specific ACRs and potential peanut-conserved were typically present at low counts, with values only slightly above zero per orthogroup (**Fig. S18**). Most other ACR categories were close to zero per orthogroup across the majority of orthogroups, largely independent of expression patterns (**Fig. S18**). Consequently, direct comparisons based on raw ACR counts are strongly influenced by background differences in ACR abundance and may obscure biologically meaningful associations. In addition, it is important to note that ACRs do not necessarily lead to gene up-regulation; increased chromatin accessibility can also be associated with gene repression (2). We therefore used enrichment-based analyses, which more effectively capture relative overrepresentation while minimizing confounding effects of background ACR abundance.

Because distal ACRs are more likely to be incorrectly assigned to targeted genes, we performed an additional enrichment analysis excluding distal ACRs. Both the full and filtered datasets showed comparable enrichment patterns, and overall trends were consistent between leaf and root tissues (**Fig. S19** and **Table S21**). Although few categories reached statistical significance, the direction and magnitude of Z-scores revealed consistent and informative trends (**Fig. S19** and **Table S21**). Given current statistical power and multiple testing restrictions, we interpret these associations as trend-level and hypothesis-generating rather than definitive. Genes showing no-bias expression were positively associated with peanut-conserved:no bias, whereas biased expression categories tended to show negative enrichment, particularly parental-legacy bias expression (**Fig. 6C** and **Table S21**). In contrast, peanut-conserved:novel bias were positively enriched for genes with novel-bias expression, but negatively or weakly enriched for no-bias and parental-legacy bias expression (**Fig.6C** and **Table S21**). Subgenome-specific ACRs exhibited similar tendencies, displaying negative enrichment with no-bias expression but positive enrichment with novel-bias and parental-legacy bias expression (**Fig. 6C** and **Table S21**). Lineage-specific ACRs and peanut-conserved:parental-legacy bias were generally positively enriched across all expression categories, with lineage-specific ACRs showing the strongest and statistically significant association with parental-legacy bias expression (**Fig. 6C** and **Table S21**). Conversely, polyploid-specific ACRs displayed consistently negative enrichment across all expression classes, particularly for no-bias expression (**Fig. 6C** and **Table S21**).

Together, these results indicate that genes with stable, unbiased expression tend to associate with conserved regulatory elements maintaining no-bias chromatin accessibility (peanut-conserved:no bias). In contrast, genes with biased expression shows enrichment for more recently evolved or divergent ACRs, such as lineage-specific ACRs and subgenome-specific ACRs that have experienced sequence gain and loss across different evolutionary trajectories, as well as peanut-conserved:novel bias ACRs, which exhibit chromatin accessibility despite high sequence similarity. Meanwhile, the correspondence between ACR evolutionary origin and expression biases, where parental-legacy bias expression associates with lineage-specific ACRs and novel bias expression aligns with subgenome-specific ACRs, reveals a coordinated regulatory evolution, highlighting how ACRs with different evolutionary histories contribute to expression asymmetry in polyploid genome.

## Discussion

Genetic variation within *cis*-regulatory elements is a major source of phenotypic diversity and evolutionary innovation. Understanding its evolutionary dynamics is therefore essential for elucidating the genetic basis of natural and artificial selection, and for facilitating targeted manipulation of gene regulation (6, 10, 56, 57). Although regulatory sequences exhibit rapid turnover across distantly related plant lineages (12), deeply conserved noncoding elements have also persisted across species diverged over 300 million years (10). However, how *cis*-regulatory elements evolve among closely related species and on lineage-specific timescale remains poorly understood (10, 12–14). Here we address this gap by systematically examining the evolutionary trajectories of regulatory regions in allotetraploid peanut and its diploid progenitors (**Fig. 2A**).

Beyond associating genetic variation with expression divergence in polyploids (16–19, 22, 58), we further investigated the evolutionary origins of regulatory variation, offering an additional dimension to understand *cis*-regulatory dynamics underlying expression changes. We uncovered a strong correspondence between homoeolog expression bias and ACR variation at both sequence and accessibility levels, shaped by ACRs arising from different evolutionary trajectories. For example, parental-legacy bias expression was enriched for lineage-specific ACRs, that inherited from corresponding progenitor; whereas novel-bias expression tends to associate with subgenome-specific, potentially *de novo* arising from subgenome or peanut-conserved:novel-bias ACRs, that with high sequence similarity but varied chromatin accessibility (**Fig. 6C** and **Table S21**). Notably, *cis*-regulatory elements typically function as components of *cis*-regulatory modules that can exert activating or repressing effects on target gene expression, such as enhancers and silencers (2). In addition, homoeolog expression bias could also be influenced by other factors, including TEs and trans-regulatory effects (59). Future investigations are needed to disentangle the individual and combined contributions of these factors to expression bias. We also detected a pronounced parental-legacy effect on regulatory evolution (17, 45, 60–62), supported by lineage-specific ACRs (**Fig. 2B**), peanut-conserved:parental-legacy bias ACRs (**Fig. 3C**) and conservation of CNSs between a subgenome and its corresponding progenitor (**Fig. 4D-4E**). However, how the strength of this parental-legacy effect shifts over time remains an important open question. Collectively, these findings highlight that ACRs with distinct evolutionary trajectories exert differential impacts on regulatory landscapes in the polyploid genome.

From a broader evolutionary perspective, we characterized the fates of regulatory regions. Most ACRs were sequence-conserved (**Fig. 2B**), exhibiting exceptionally high sequence similarity (mean BLASTed ratio ∼0.98 and mean mismatch rate ∼0.02) (**Fig. 2D**), despite the ∼2 Mya divergence between the diploid progenitors (28). Many of these sequence-conserved ACRs also maintained comparable histone modification profiles (**Fig. 2E**) and chromatin accessibility (**Fig. 3C**), likely reflecting both the recent formation of the polyploid and strong selective constrains on regulatory function (10, 49, 50). Meanwhile, we also observed the emergence of new ACRs that originated either *de novo* from previously non-regulatory regions or through accumulated mutations in preexisting regulatory sequences (6, 9–11). Polyploid-specific or subgenome-specific ACRs exemplify *de novo* origins, arising after hybridization or after polyploidization (**Fig. 2A**). TEs might contribute substantially to these *de novo* origins, notably, more than half of subgenome-specific ACRs overlap with TEs (**Fig. 2C** and **Fig. S5**). It is also possible that these polyploid- or subgenome-specific ACRs result from loss in the progenitor genomes, but the likelihood of simultaneous loss across multiple progenitors should be low. In contrast, some lineage-specific ACRs likely represent gradual sequence divergence: ACRs that shift from peanut-conserved to lineage-specific when the BLAST similarity threshold increases from 0.5 to 0.8 (**Fig. 2B**) illustrate this process: these regions likely existed prior to the divergence of the diploid progenitors and subsequently accumulated mutations, increasing intra-lineage similarity to ∼80% while reducing inter-lineage similarity to ∼50%. Peanut-conserved ACRs with high sequence similarity but biased chromatin accessibility provide another example of diversification (**Fig. 5**). The generally lower frequency of subgenome- or polyploid-specific ACRs, compared with the more abundant lineage-specific ACRs or peanut-conserved ACRs with biased chromatin accessibility, supports the conclusion that regulatory diversification through gradual mutation accumulation is more common than *de novo* emergence (10, 11, 22).

Importantly, we found that even minor sequence changes can have disproportionately large effects on regulatory activity. Peanut-conserved ACRs with biased chromatin accessibility exemplify this, where they exhibited high sequence similarity but substantial chromatin accessibility differences that often manifest as presence/absence patterns rather than gradual quantitative changes (**Fig. 3D**). Furthermore, we demonstrate a correlation between changes in CNS composition and chromatin accessibility bias, a relationship previously not often examined in lineage-specific contexts (10, 63). For instance, the peanut-conserved:parental-legacy bias ACRs always possessed more CNSs that conserved between the respective subgenome and its corresponding diploid, whereas peanut-conserved:no bias ACRs were enriched for CNSs that conserved across four genomes (**Fig. 5**). Although these associations are compelling, they remain correlative. Without allele-specific assays or targeted perturbations, we cannot ascribe causality to CNS turnover, and trans-acting factors, peak-calling thresholds, or tissue composition may also contribute (64–66). A limitation is inherent to CNS identification itself, which requires a predefined degree of sequence conservation. The large number of CNSs conserved exclusively in one or both progenitors but absent from the subgenomes (**Fig. 4D-4E**) may suggest that these sequences have undergone functional divergence in the subgenomes, shifting toward new regulatory roles at the expense of detectable sequence conservation, even if the underlying sequence changes might be minor. On the other hand, examining transcription factor binding motifs (67) or footprinting data (56, 68) will also be essential for understanding how conserved elements transition into novel regulatory functions and how frequently such events occur.

In summary, our study provides empirical evidence illuminating the evolutionary dynamics of regulatory regions in polyploid plants. By integrating sequence variation, CNSs turnover, chromatin accessibility, histone modification, and expression divergence, we provide a comprehensive view of how *cis*-regulatory evolution drives expression diversification and contributes to evolutionary innovation in plant genome.

## Materials and Methods

### Plant materials

Three Arachis species were used in this study: the cultivated allotetraploid peanut *A. hypogaea* and its two diploid progenitors, *A. duranensis* and *A. ipaensis*. Plants were grown in soil for approximately 10 days at 25 °C under photoperiodic lighting (16 h light–8 h dark). Mature leaf and root tissues were collected with two biological replicates per tissue for each species for library preparation. Details of library preparation and data preprocessing are provided in the supplementary materials.

### Sequence conservation of ACRs identified in polyploidy peanut genome

To investigate the sequence ancestry of ACRs identified in the two subgenomes of polyploid peanut, we adopted a BLAST-based approach following the methods described in (13). ACR sequences were used as queries, and reference regions were defined as the genomic intervals spanning the two orthologous genes flanking each ACR in the corresponding genomes (Fig. S2). These reference regions were identified using GENESPACE (69) and analyses were restricted to regions conserved across all four genomes.

ACRs from each subgenome were BLASTed against the reference regions using BLAST (70) using the same parameters as described in (13). The resulting BLAST outputs were then combined and filtered using customized scripts. Sequence divergence was quantified using two metrics: the BLASTed ratio and the mismatch rate (22). Differences in these metrics among the four ACR groups, lineage-specific, peanut-conserved, potential peanut-conserved and polyploid-specific ACRs, were assessed using ANOVA and followed by Games-Howell *post hoc* tests to account for heterogeneity of variances.

### Differential chromatin accessibility

To examine quantitative differences in ACRs between the two tissue types from the same species, we generated four pseudo replicates by randomly sampling the equal number of clean reads from the two biological replicates using a customized Python script. Normalized read counts were then calculated for each peak, and DESeq2 (71) was used to identify regions with significant differences in accessibility. Peaks with an adjusted P-value (FDR) < 0.05 and an absolute log_2_ fold change > 1 were defined as differential ACRs.

The same analytical framework was used to examine accessibility bias across peanut-conserved, lineage-specific and polyploid-specific ACRs and their corresponding BLASTed regions in the reference genomes (**Fig. 2B**). Parental no bias was defined as the absence of significant accessibility differences between the two diploids and between the two subgenomes. Parental subgenome A or B-biased accessibility was assigned when a subgenome and its corresponding progenitor both exhibited significantly higher accessibility relative to the other pair. No bias in progeny referred to cases with no significant difference between the two subgenomes, further divided into progenitor A- or B-biased when one progenitor showed significantly higher chromatin accessibility. Novel bias in progeny was defined as subgenome A- or B- biased accessibility in the absence of significant differences between the progenitors. Finally, reverse bias in progeny (subgenome A- or subgenome B-biased) referred to cases where a subgenome showed significantly higher chromatin accessibility, but its corresponding progenitor had significantly lower chromatin accessibility.

### Analyses of CNS dynamics within polyploid-diploid framework

CNS datasets of three peanut species, *A. hypogaea, A. duranensis* and *A. ipaensis* were downloaded from Conservatory (www.conservatorycns.com). We summarized the counts, repeats, and conservation states for each CNS across the three peanut species or four genomes using customed R script. When examining the CNSs located within the sequence-conserved ACRs, we first used BEDtools (72) to intersect the coordinates of these CNSs with the coordinates of sequence-conserved ACRs or their corresponding BLASTed regions, with the requirement that CNSs might be completely within the ACRs or BLASTed regions. Then we analyzed the CNS conservation states within each ACR and its corresponding BLASTed region, requiring CNSs associated with genes belonging to the same orthogroup.

To explore the relative enrichment of distinct CNS conservation states across varied chromatin accessibility of sequence-conserved ACRs, we generated a rank-normalized matrix from the CNS conservation state count matrix. For each CNS, we first identified the states with non-zero counts. States with zero counts were treated as absent and assigned a rank-normalized score of zero. For states with positive values, we computed ranks based on descending abundance, such that the highest count received rank 1 and the lowest received rank N, where N is the number of states present. These ranks were then normalized by dividing by N, producing values between 1/N and 1. The resulting rank-normalized scores were placed back into a full vector, with zeros retained for absent states. This procedure was applied row-wise to the CNS matrix, producing a final rank-normalized matrix with the same dimensions as the original input.

### Enrichment between expression bias and ACR types

Only Orthogroups exclusively associated with ACRs that had been assigned an evolutionary origin were included in the analysis. Expression bias categories were simplified into four types: no bias (including “no bias in progeny” and “parental no bias”), novel bias (including “novel bias” and “novel bias reverse”), parental-legacy bias (including “parental subgenome A-biased” and “parental subgenome B-biased”), and others. Associated ACRs were first classified by sequence conservation and then by chromatin accessibility bias patterns. Permutation-based enrichment analyses were performed separately for leaf and root tissues using both unfiltered and filtered datasets (with distal ACRs removed).

For each Orthogroup, the proportion of ACRs belonging to each category was calculated by dividing the number of ACRs of that type by the total number of ACRs within that Orthogroup. These normalized proportions were then aggregated by expression bias groups. Observed mean proportions for each ACR category were calculated across Orthogroups within each expression group. To evaluate the significance of these observed proportions, 10,000 random permutations were conducted by shuffling expression bias group labels while preserving the ACR composition within each Orthogroup. For each permutation, mean proportions were recalculated to generate a null distribution. The null mean and standard deviation were then used to compute standardized Z-scores, and empirical P-values were derived as the proportion of permuted means equal to or greater than the observed mean proportion. Multiple testing correction was applied using the Benjamini–Hochberg (BH) method, and categories with adjusted P < 0.05 were considered significantly enriched.

### Data, Materials and Software Availability

All the raw reads of ATAC-seq, ChIP-seq and RNA were uploaded to the National Center for Biotechnology Information (NCBI) under the BioProject accession PRJNA1373159. Data analysis scripts and relevant raw data can be found at: https://github.com/Xianglichina/peanut_atac_blast.

## Acknowledgments

We sincerely appreciate the members of the Schmitz lab for their valuable analytical suggestions. J.P.M. was supported by an NIH Training Grant (T32GM142623). This research was supported by the National Science Foundation (IOS-1856627) awarded to R.J.S.

## References

1. A. P. Marand, A. L. Eveland, K. Kaufmann, N. M. Springer, cis-Regulatory Elements in Plant Development, Adaptation, and Evolution. Annu. Rev. Plant Biol. 74, 111–137 (2023).

2. R. J. Schmitz, E. Grotewold, M. Stam, Cis-regulatory sequences in plants: Their importance, discovery, and future challenges. Plant Cell 34, 718–741 (2022).

3. S. L. Klemm, Z. Shipony, W. J. Greenleaf, Chromatin accessibility and the regulatory epigenome. Nat. Rev. Genet. 20, 207–220 (2019).

4. P. J. Wittkopp, G. Kalay, Cis-regulatory elements: molecular mechanisms and evolutionary processes underlying divergence. Nature Reviews Genetics 13, 59–69 (2012).

5. H. K. Long, S. L. Prescott, J. Wysocka, Ever-Changing Landscapes: Transcriptional Enhancers in Development and Evolution. Cell 167, 1170–1187 (2016).

6. X. Li, R. J. Schmitz, Cis-regulatory dynamics in plant domestication. Trends Genet. 0 (2025).

7. P. S. Soltis, D. E. Soltis, Plant genomes: Markers of evolutionary history and drivers of evolutionary change. Plants People Planet 3, 74–82 (2021).

8. F. Murat, Y. Van de Peer, J. Salse, Decoding plant and animal genome plasticity from differential paleo-evolutionary patterns and processes. Genome Biol. Evol. 4, 917–928 (2012).

9. A. Katikaneni, C. B. Lowe, Novelty versus innovation of gene regulatory elements in human evolution and disease. Curr. Opin. Genet. Dev. 90, 102279 (2025).

10. K. R. Amundson, et al., A deep-time landscape of plant cis-regulatory sequence evolution. bioRxiv 2025.09.17.676453 (2025).

11. J. M. C. McDonald, R. D. Reed, Beyond modular enhancers: new questions in cis-regulatory evolution. Trends Ecol. Evol. 0 (2024).

12. Z. Lu, et al., The prevalence, evolution and chromatin signatures of plant regulatory elements. Nat Plants 5, 1250–1259 (2019).

13. J. P. Mendieta, et al., Investigating the cis-regulatory basis of C3 and C4 photosynthesis in grasses at single-cell resolution. Proc. Natl. Acad. Sci. U. S. A. 121, e2402781121 (2024).

14. H. Yan, et al., A single-cell rice atlas integrates multi-species data to reveal cis-regulatory evolution. Nat. Plants 11, 2050–2071 (2025).

15. E. Kuzmin, J. S. Taylor, C. Boone, Retention of duplicated genes in evolution. Trends Genet. 38, 59–72 (2022).

16. C. Fang, et al., Dynamics of cis-regulatory sequences and transcriptional divergence of duplicated genes in soybean. Proc. Natl. Acad. Sci. U. S. A. 120, e2303836120 (2023).

17. G. Hu, et al., Evolutionary Dynamics of Chromatin Structure and Duplicate Gene Expression in Diploid and Allopolyploid Cotton. Mol. Biol. Evol. 41 (2024).

18. J. Han, et al., Genome-wide chromatin accessibility analysis unveils open chromatin convergent evolution during polyploidization in cotton. Proc. Natl. Acad. Sci. U. S. A. 119, e2209743119 (2022).

19. C. Fang, et al., Dynamics of accessible chromatin regions and subgenome dominance in octoploid strawberry. Nat. Commun. 15, 1–14 (2024).

20. F. Almeida-Silva, Y. Van de Peer, Gene expression divergence following gene and genome duplications in spatially resolved plant transcriptomes. Plant Cell 37, koaf243 (2025).

21. T. Srikant, A. Gonzalo, K. Bomblies, Chromatin accessibility and gene expression vary between a new and evolved autopolyploid of Arabidopsis arenosa. Mol. Biol. Evol. 41, msae213 (2024).

22. X. Li, X. Zhang, R. J. Schmitz, From duplication to divergence: single-cell insights into transcriptional and cis-regulatory landscapes in soybean. Plant Cell koaf279 (2025).

23. G. Seijo, et al., Genomic relationships between the cultivated peanut (Arachis hypogaea, Leguminosae) and its close relatives revealed by double GISH. Am. J. Bot. 94, 1963–1971 (2007).

24. G. Robledo, G. I. Lavia, G. Seijo, Species relations among wild Arachis species with the A genome as revealed by FISH mapping of rDNA loci and heterochromatin detection. Theor. Appl. Genet. 118, 1295–1307 (2009).

25. D. J. Bertioli, et al., The genome sequences of Arachis duranensis and Arachis ipaensis, the diploid ancestors of cultivated peanut. Nat. Genet. 48, 438–446 (2016).

26. X. Chen, et al., Draft genome of the peanut A-genome progenitor (Arachis duranensis) provides insights into geocarpy, oil biosynthesis, and allergens. Proc. Natl. Acad. Sci. U. S. A. 113, 6785–6790 (2016).

27. Q. Lu, et al., Genome Sequencing and Analysis of the Peanut B-Genome Progenitor (Arachis ipaensis). Front. Plant Sci. 9, 604 (2018).

28. X. Chen, et al., Sequencing of Cultivated Peanut, Arachis hypogaea, Yields Insights into Genome Evolution and Oil Improvement. Mol. Plant 12, 920–934 (2019).

29. K. Zhao, et al., Pangenome analysis reveals structural variation associated with seed size and weight traits in peanut. Nat. Genet. 1–12 (2025).

30. X. Zhang, et al., A spatially resolved multi-omic single-cell atlas of soybean development. Cell 188, 550–567.e19 (2024).

31. A. Farmer, S. Thibivilliers, K. H. Ryu, J. Schiefelbein, M. Libault, Single-nucleus RNA and ATAC sequencing reveals the impact of chromatin accessibility on gene expression in Arabidopsis roots at the single-cell level. Mol. Plant 14, 372–383 (2021).

32. M. W. Dorrity, et al., The regulatory landscape of Arabidopsis thaliana roots at single-cell resolution. Nat. Commun. 12, 3334 (2021).

33. V. Sundaram, J. Wysocka, Transposable elements as a potent source of diverse cis-regulatory sequences in mammalian genomes. Philos. Trans. R. Soc. Lond. B Biol. Sci. 375, 20190347 (2020).

34. A. Y. Du, J. D. Chobirko, X. Zhuo, C. Feschotte, T. Wang, Regulatory transposable elements in the encyclopedia of DNA elements. Nat. Commun. 15, 7594 (2024).

35. W. A. Ricci, et al., Widespread long-range cis-regulatory elements in the maize genome. Nat Plants (2019).

36. A. E. Yocca, Z. Lu, R. J. Schmitz, M. Freeling, P. P. Edger, Evolution of Conserved Noncoding Sequences in Arabidopsis thaliana. Mol. Biol. Evol. 38, 2692–2703 (2021).

37. L. Quadrana, I. R. Henderson, The natural history of transposons in plant pangenomes and panepigenomes. Curr. Opin. Plant Biol. 88, 102818 (2025).

38. S. Moser Tralamazza, L. Nanchira Abraham, C. S. Reyes-Avila, B. Corrêa, D. Croll, Histone H3K27 methylation perturbs transcriptional robustness and underpins dispensability of highly conserved genes in fungi. Mol. Biol. Evol. 39, msab323 (2022).

39. G. Vernaz, et al., Epigenetic divergence during early stages of speciation in an African crater lake cichlid fish. Nat. Ecol. Evol. 6, 1940–1951 (2022).

40. J. P. Mendieta, A. P. Marand, W. A. Ricci, X. Zhang, R. J. Schmitz, Leveraging histone modifications to improve genome annotations. G3 11 (2021).

41. H. Le, C. H. Simmons, X. Zhong, Functions and mechanisms of histone modifications in plants. Annu. Rev. Plant Biol. 76, 551–578 (2025).

42. X. Zhang, Y. V. Bernatavichute, S. Cokus, M. Pellegrini, S. E. Jacobsen, Genome-wide analysis of mono-, di- and trimethylation of histone H3 lysine 4 in Arabidopsis thaliana. Genome Biol. 10, R62 (2009).

43. F. Roudier, et al., Integrative epigenomic mapping defines four main chromatin states in Arabidopsis: Organization of the Arabidopsis epigenome. EMBO J. 30, 1928–1938 (2011).

44. H. Yu, et al., Epigenetic insights into the domestication of tetraploid peanut. Plant Physiol. 198, kiaf254 (2025).

45. M.-J. Yoo, E. Szadkowski, J. F. Wendel, Homoeolog expression bias and expression level dominance in allopolyploid cotton. Heredity (Edinb.) 110, 171–180 (2013).

46. J. Wu, et al., Homoeolog expression bias and expression level dominance in resynthesized allopolyploid Brassica napus. BMC Genomics 19, 586 (2018).

47. C. E. Grover, et al., Homoeolog expression bias and expression level dominance in allopolyploids. New Phytol. 196, 966–971 (2012).

48. M. Li, R. Wang, X. Wu, J. Wang, Homoeolog expression bias and expression level dominance (ELD) in four tissues of natural allotetraploid Brassica napus. BMC Genomics 21, 330 (2020).

49. A. Hendelman, et al., Conserved pleiotropy of an ancient plant homeobox gene uncovered by cis-regulatory dissection. Cell (2021).

50. A. Lanctot, A. Hendelman, P. Udilovich, G. M. Robitaille, Z. B. Lippman, Antagonizing cis-regulatory elements of a conserved flowering gene mediate developmental robustness. Proc. Natl. Acad. Sci. U. S. A. 122, e2421990122 (2025).

51. D. Ciren, S. Zebell, Z. B. Lippman, Extreme restructuring of cis-regulatory regions controlling a deeply conserved plant stem cell regulator. PLoS Genet. 20, e1011174 (2024).

52. X. Wang, et al., Dissecting cis-regulatory control of quantitative trait variation in a plant stem cell circuit. Nat Plants 7, 419–427 (2021).

53. T. C. Tran, et al., Altered interactions between cis-regulatory elements partially resolve BLADE-ON-PETIOLE genetic redundancy in Capsella rubella. Plant Cell 36, 4637–4657 (2024).

54. Y. Luo, et al., Characterization and functional analysis of conserved non-coding sequences among poaceae: insights into gene regulation and phenotypic variation in maize. BMC Genomics 26, 46 (2025).

55. J. Van de Velde, M. Van Bel, D. Vaneechoutte, K. Vandepoele, A collection of conserved noncoding sequences to study gene regulation in flowering plants. Plant Physiol. 171, 2586–2598 (2016).

56. A. P. Marand, et al., The genetic architecture of cell type-specific cis regulation in maize. Science 388, eads6601 (2025).

57. D. Villar, P. Flicek, D. T. Odom, Evolution of transcription factor binding in metazoans - mechanisms and functional implications. Nat. Rev. Genet. 15, 221–233 (2014).

58. M. Li, W. Sun, F. Wang, X. Wu, J. Wang, Asymmetric epigenetic modification and homoeolog expression bias in the establishment and evolution of allopolyploid Brassica napus. New Phytol. 232, 898–913 (2021).

59. S. Bottani, N. R. Zabet, J. F. Wendel, R. A. Veitia, Gene Expression Dominance in Allopolyploids: Hypotheses and Models. Trends Plant Sci. 23, 393–402 (2018).

60. K. A. Bird, et al., Replaying the evolutionary tape to investigate subgenome dominance in allopolyploid Brassica napus. New Phytol. (2020).

61. P. P. Edger, et al., Origin and evolution of the octoploid strawberry genome. Nat. Genet. 51, 541–547 (2019).

62. R. J. A. Buggs, et al., The legacy of diploid progenitors in allopolyploid gene expression patterns. Philos. Trans. R. Soc. Lond. B Biol. Sci. 369, 20130354 (2014).

63. J. C. Schnable, B. S. Pedersen, S. Subramaniam, M. Freeling, Dose-sensitivity, conserved non-coding sequences, and duplicate gene retention through multiple tetraploidies in the grasses. Front. Plant Sci. 2, 2 (2011).

64. M. A. A. Minow, et al., Hybrids reveal accessible chromatin trans genetic associations. bioRxivorg 2025.10.09.681407 (2025).

65. M. A. Ballinger, K. L. Mack, S. M. Durkin, E. A. Riddell, M. W. Nachman, Environmentally robust cis-regulatory changes underlie rapid climatic adaptation. Proc. Natl. Acad. Sci. U. S. A. 120, e2214614120 (2023).

66. M. Lensink, G. Monroe, D. J. Kliebenstein, Trans-regulatory loci shape natural variation of gene expression plasticity in Arabidopsis. Genetics 230, iyaf116 (2025).

67. L. A. Baumgart, et al., Recruitment, rewiring and deep conservation in flowering plant gene regulation. Nat. Plants 11, 1514–1527 (2025).

68. Y. Hu, et al., Multiscale footprints reveal the organization of cis-regulatory elements. Nature 638, 779–786 (2025).

69. J. T. Lovell, et al., GENESPACE tracks regions of interest and gene copy number variation across multiple genomes. Elife 11 (2022).

70. C. Camacho, et al., BLAST+: architecture and applications. BMC Bioinformatics 10, 421 (2009).

71. M. I. Love, W. Huber, S. Anders, Moderated estimation of fold change and dispersion for RNA-seq data with DESeq2. Genome Biol. 15, 550 (2014).

72. A. R. Quinlan, I. M. Hall, BEDTools: a flexible suite of utilities for comparing genomic features. Bioinformatics 26, 841–842 (2010).

